# A fluorescently labelled quaternary ammonium compound (NBD-DDA) to study mode-of-action and resistance mechanisms in bacteria

**DOI:** 10.1101/2022.07.15.500178

**Authors:** Niclas Nordholt, Kate O’Hara, Ute Resch-Genger, Mark A. T. Blaskovich, Bastian Rühle, Frank Schreiber

## Abstract

Quaternary ammonium compounds (QACs) are widely used as active agents in disinfectants, antiseptics, and preservatives. Despite being in use since the 1940s, there remain multiple open questions regarding their detailed mode-of-action and the mechanisms, including phenotypic heterogeneity, that can make bacteria less susceptible to QACs. To facilitate mode-of-action studies, we synthesized a fluorescent analogue of the quaternary ammonium compound benzalkonium chloride, namely *N*-dodecyl-*N,N*-dimethyl-[2-[(4-nitro-2,1,3-benzoxadiazol-7-yl)amino]ethyl]azanium-iodide (NBD-DDA). NBD-DDA is readily detected by flow cytometry and fluorescence microscopy with standard GFP/FITC-settings, making it suitable for molecular and single-cell studies. NBD-DDA was then used to investigate resistance mechanisms which can be heterogeneous among individual bacterial cells. Our results reveal that the antimicrobial activity of NBD-DDA against *E. coli, S. aureus* and *P. aeruginosa* is comparable to that of benzalkonium chloride (BAC), a widely used QAC. Characteristic time-kill kinetics and increased tolerance of a BAC tolerant *E. coli* strain against NBD-DDA suggest that the mode of action of NBD-DDA is similar to that of BAC. Leveraging these findings and NBD-DDA’s fluorescent properties, we show that reduced cellular adsorption is responsible for the evolved BAC tolerance in the BAC tolerant *E. coli* strain. As revealed by confocal laser scanning microscopy (CLSM), NBD-DDA is preferentially localized in the cell envelope of *E. coli*, which is a primary target of BAC and other QACs. Overall, NBD-DDA’s antimicrobial activity, its fluorescent properties, and its ease of detection render it a powerful tool to study the mode-of-action and the resistance mechanisms of QACs in bacteria.

## Introduction

Quaternary ammonium compounds (QACs) were first admitted to the market in the 1940s and are widely used as disinfectants, preservatives, surfactants, and antiseptics ^1^. They are positively charged surfactants that possess bactericidal, fungicidal, and viricidal activity ^2,3^. QACs are named after a quaternary nitrogen comprising the cationic headgroup, which carries hydrophobic moieties (Figure 1). Among the most widely used QACs is benzalkonium chloride (BAC). Benzalkonium chloride is a cationic surfactant and consists of a mixture of alkyl-benzyl-dimethyl-ammonium chlorides (ADBAC) with alkyl chain lengths varying from 8 to 18 carbon atoms (Figure 1). ADBACs with the alkyl-chain lengths C_12_, C_14_, and C_16_ possess the highest antimicrobial activity ^4,5^ and typically make up the largest mass fraction in BAC ^6^.

**Figure 1:**
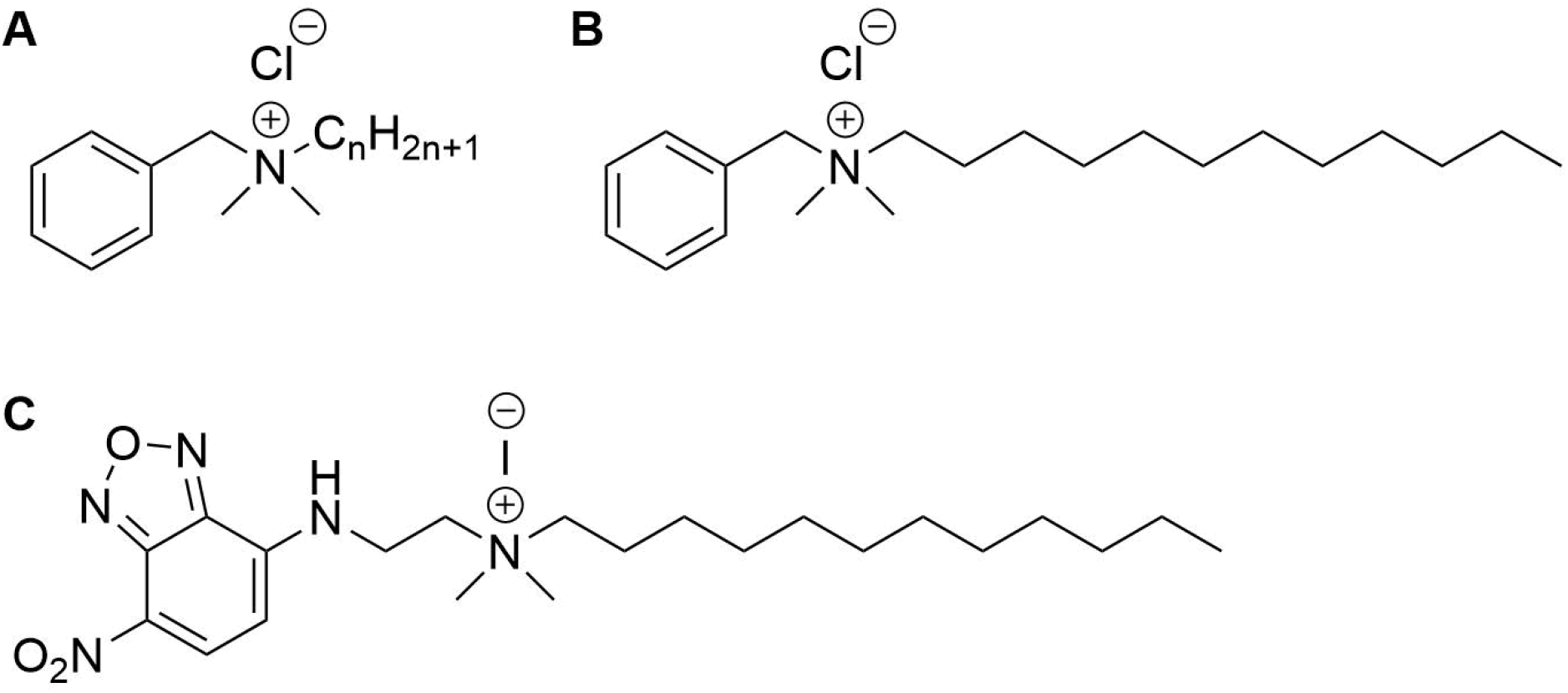
Structures of the compounds used in this study. A) Benzalkonium chloride, with n ranging from 8 to 18, B) benzyl-dimethyl-dodecylammoniumchlorid (BAC_12_) and C) N-dodecyl-N,N-dimethyl-[2-[(4-nitro-2,1,3-benzoxadiazol-7-yl)amino]ethyl]azanium-iodide (NBD-DDA).

The main target of BAC is thought to be the bacterial cell envelope ^2,7^. The general mode of action likely proceeds as follows: the cationic headgroup attaches to anionic sites on the cell surface, which possesses a net negative charge. Displacement of membrane stabilizing cations (Ca^2+^, Mg^2+^) and integration of the hydrophobic region into the membrane cause membrane instability, changes in membrane fluidity, and leakage of cytoplasmic material ^2,8^. The mode of action of BAC and other QACs appears to be dose dependent and the detailed mechanisms and their kinetics are still not fully understood ^3,8–10^.

Resistance mechanisms to BAC and other QACs involve changes to the cell envelope, multi-drug efflux, and target degradation, as well as increased biofilm formation ^1,11^. The fact that there are still genes being identified that play a role in the reduced susceptibility to BAC suggests that the exact mechanisms and pathways that reduce the susceptibility to BAC and other QACs remain underexplored ^12^. For many mutations that confer resistance or tolerance to BAC and related compounds, the mechanistic consequence for resistance is not known. For example, do mutations in genes pertaining the cell envelope reduce the rate or amount of BAC adsorbed to the outer membrane? Or do the changes in membrane composition increase membrane stability and, thus, allow bacteria to adsorb larger amounts of BAC before the membrane is compromised? How does efflux contribute to reduced susceptibility? How does BAC interact with components of the periplasm and the cytoplasm? And what are the kinetics of BAC induced death, from adsorption of BAC to cell death? Another open question is to which extent phenotypic heterogeneity affects the survival of exposure to BAC, and what are the determinants of individual cells that make them more or less susceptible to BAC.

In this study, we synthesized a fluorescent BAC analogue, NBD-DDA, providing a valuable tool to address open research questions related to BAC and similar QACs. We characterized its antimicrobial activity against three bacterial species relevant for antimicrobial resistance (AMR), *E. coli, S. aureus*, and *P. aeruginosa*. Furthermore, we demonstrated the potential of NBD-DDA for single-cell studies with fluorescence microscopy and flow cytometry. In the future, we anticipate NBD-DDA to become a powerful tool to advance the knowledge on the mode-of-action and reaction kinetics of BAC and related QACs with bacteria and to study the mechanisms that govern the reduced susceptibility of microorganisms to these compounds.

## Results and Discussion

### Synthesis of NBD-DDA and chemical characterization

Benzalkonium chlorides (*N*-Alkyl-*N*-benzyl-*N,N*-dimethylammonium chlorides; BAC) are a class of compounds with the general structure depicted in Figure 1 A. When employed as biocides, often a mixture of BAC with different alkyl chain lengths are used, with compounds with C_12_, C_14_, and C_16_ chains possessing the highest antimicrobial activity ^4,5^ and typically making up the largest mass fraction in BAC biocides ^6^. Fluorophore-labeled derivatives such as NBD-DDA (Figure 1 C) can be readily synthesized in two steps from commercially available precursors. In the first step, *N,N*-dimethylethylenediamine is reacted with NBD-chloride in a nucleophilic substitution (S_N_) reaction, yielding the intermediate product *N,N*-dimethyl-*N*’-(4-nitro-5-benzofurazanyl)-1,2-ethanediamine (NBD-DMA) ^13^. This can then be reacted with alkyl halides (especially alkyl iodides) of various chain lengths in another S_N_ reaction, yielding the final NBD labeled quaternary ammonium compound.

In this study, we chose dodecyl iodide for the functionalization in the second step to obtain a derivative (NBD-DDA) that mimics one of the most active compounds typically found in BAC mixtures (Figure 1 B, C). Importantly, compounds similar to NBD-DDA with varying alkyl-chain lengths mimicking BAC components other than the C_12_ derivative or even using benzyl or aryl groups instead of alkyl groups can be synthesized with minor modifications to the synthesis protocol. This can pave the way for mechanistic studies of the antimicrobial activity of different alkyl-chain lengths in BACs, which can vary between bacterial species ^4,5^.

The dye NBD was chosen due to its small size, its structural similarity to a simple aromatic substituent and its fluorescence emission in the visible spectrum, which can be excited with typical laser or light emitting diode (LED) light sources around 488 nm. Another advantage of NBD is the insensitivity of its optical properties towards changes in chloride concentration and pH in the physiologically relevant range of pH 5.0-7.4. Moreover, NBD dyes are popular lipid membrane probes owing to the ability of their derivatives to be readily incorporated into lipid membranes and the sensitivity of their optical properties to the polarity of the dye microenvironment ^13–15^.

The chemical structure and purity of NBD-DDA was confirmed by high resolution mass spectrometry (HR-MS) and NMR spectroscopy (Figure 2). Subsequently, the absorption and emission properties of NBD-DDA were determined to clarify whether the standard FITC settings could be used for the fluorescence microscopy and flow cytometry studies. The optical characterization of the compound by fluorescence spectroscopy (Figure 3) in DMSO and PBS showed two absorption bands at 335 nm and 468 nm respectively, with the strongest absorption band being located at 468 nm. The fluorescence emission spectra show a strong emission band with a maximum at 537 nm in DMSO and a weaker fluorescence band at 542 nm in PBS. These excitation and emission characteristics are very similar to GFP and FITC, making this dye well suited for fluorescence microscopy imaging studies using the same settings as applied for GFP and FITC.

**Figure 2:**
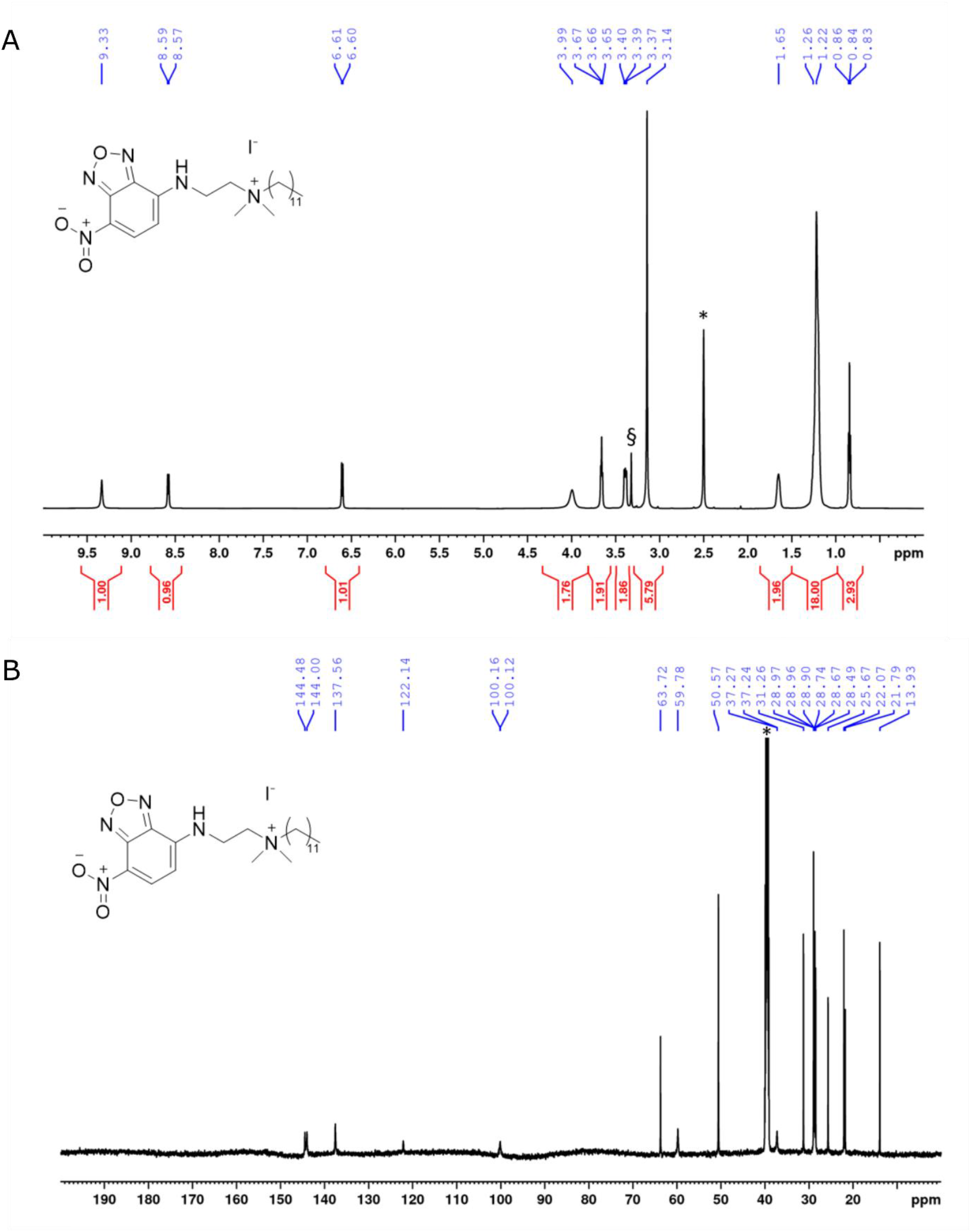
NMR spectra of NBD-DDA. A) ^1^H-NMR of NBD-DDA. * and § denote residual solvent peaks from DMSO and H_2_O, respectively. B) _13_C-NMR of NBD-DDA. * denotes the residual solvent peak from DMSO.

**Figure 3:**
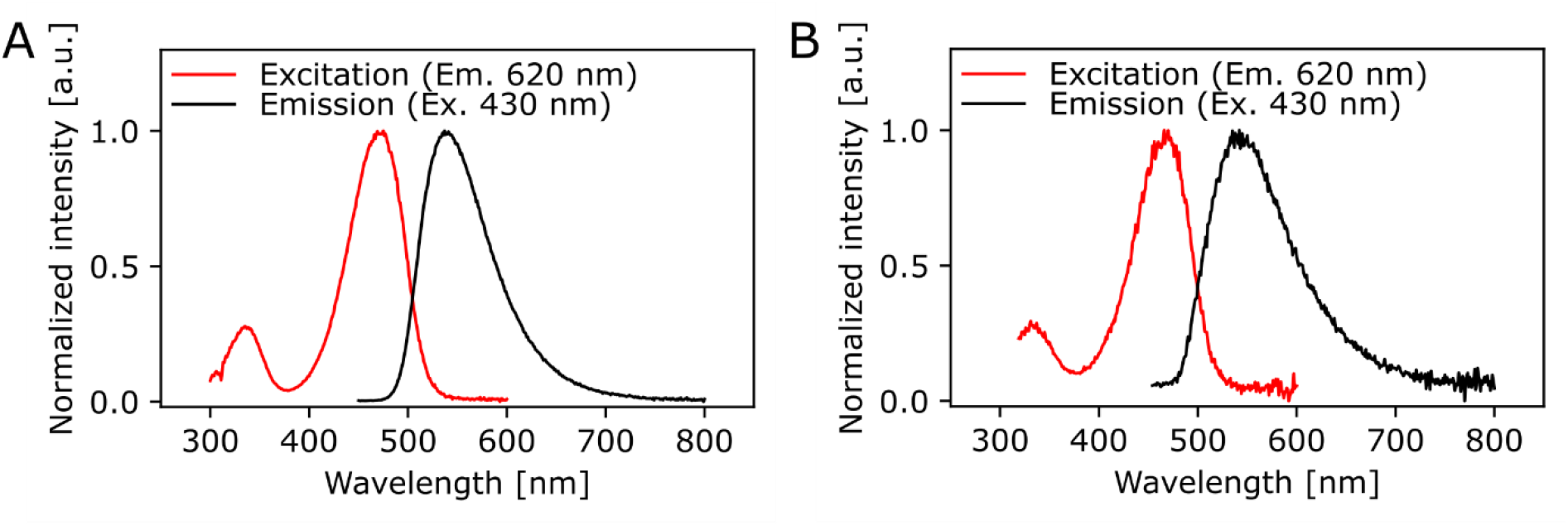
Spectrally corrected normalized fluorescence excitation and emission spectra of NBD-DDA in A) DMSO and B) PBS (1x, pH 7.4). The excitation spectra were monitored at an emission wavelength of 620 nm and the emission spectra were excited at 430 nm.

### Antimicrobial activity of NBD-DDA is comparable to BAC

The antimicrobial activity of NBD-DDA against *E. coli, S. aureus*, and *P. aeruginosa* was assessed and compared to that of BAC and benzyl-dimethyl-dodecylammonium chloride (BAC_12_). To this end, the minimum inhibitory concentration (MIC) and the minimum bactericidal concentration (MBC) were determined. The MIC is the concentration which inhibits growth within 24 hours and the MBC is the concentration which reduces the number of cfu by a factor 10^3^ or more, i.e., kills ≥ 99.9% of the cells, within 24 hours. Additionally, a BAC tolerant *E. coli* strain (*E. coli* S4) was included in the assays to test whether NBD-DDA and BAC share a similar mode of action. The strain *E. coli* S4 was isolated from a laboratory evolution experiment and shows increased short-term survival in the presence of BAC, without changes in the MIC or MBC ^12^. The MIC and MBC of all substances tested for *P. aeruginosa* were above 150 µM, the highest concentration in the assay. The MIC and MBC of NBD-DDA for all other tested strains were comparable to the MIC and MBC of BAC and BAC_12_ (Table 1). These results indicate that the fluorescent NBD moiety does not affect the antimicrobial properties of NBD-DDA.

**Table 1:** MIC and MBC values of NBD-DDA as compared to those of BAC and BAC_12_ for different bacterial strains.

| Strain | MIC* |  |  | MBC* |  |  | Medium |
| --- | --- | --- | --- | --- | --- | --- | --- |
|  | NBD-DDA | BAC<sub>12</sub> | BAC | NBD-DDA | BAC<sub>12</sub> | BAC |  |
| *E. coli* | 20 | 30 | 20 | 20 | 30 | 20 | M9 |
| *E. coli* S4 | 20 | 30 | 20 | 20 | 30 | 20 | M9 |
| *P. aeruginosa* | >150 | >150 | >150 | >150 | >150 | >150 | MH2 |
| *S. aureus* | 10 | 10 | 20 | 30 | 30 | 20 | MH2 |

\*Tested concentrations: 10, 20, 30, 40, 60, 100, and 150 μM.

### Common mode of action in BAC and NBD-DDA

In a recent study, we showed that *E. coli* populations display heterogeneity in their ability to survive lethal doses of BAC, as was apparent from bimodal killing kinetics ^12^. Here, we reasoned that if NBD-DDA has a similar mode-of-action as BAC, *E. coli* should also display bimodal killing kinetics. Indeed, this was the case when *E. coli* was exposed to a lethal concentration of NBD-DDA (Figure 4 A). These results were further corroborated by the fact that the BAC-tolerant *E. coli* strain S4 showed an increased short-term survival in the presence of NBD-DDA in comparison to the wild type (Figure 4 B). The time-kill experiments with NBD-DDA and BAC suggested a slight increase in potency of NBD-DDA as compared to BAC as killing with 30 µM NBD-DDA resulted in a comparable reduction in cfu as 60 µM BAC within 20 minutes of exposure (Figure 4 B).

**Figure 4:**
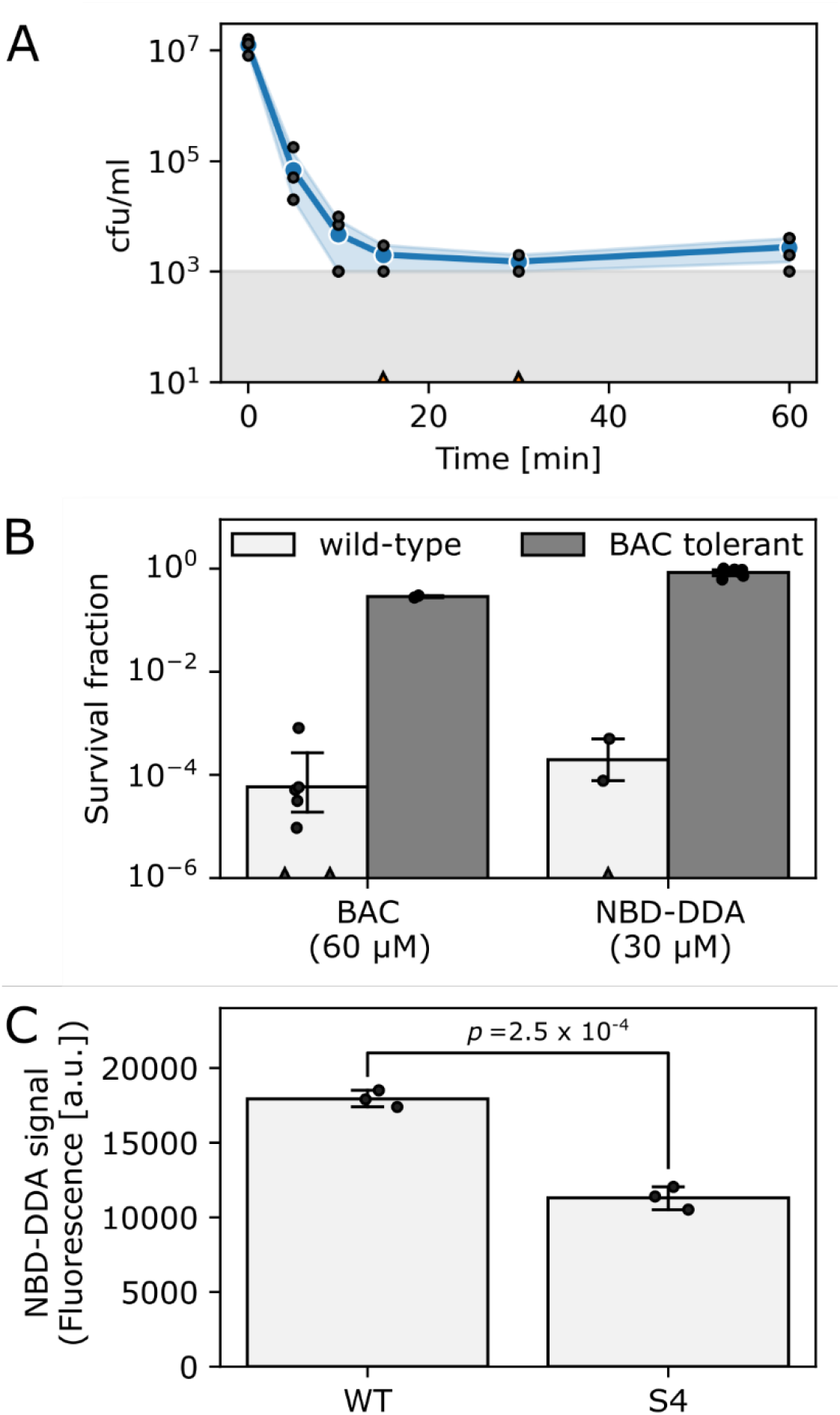
NBD-DDA and BAC share a similar mode-of-action. A) Time-kill curve of *E. coli* wild-type exposed to 30 µM NBD-DDA. B) Survival of *E. coli* wild-type and the BAC tolerant *E. coli* S4 after 20 minutes of treatment with BAC or NBD-DDA. C) NBD-DDA fluorescence is reduced in the BAC tolerant strain *E. coli* S4, suggesting that reduced adsorption as underlying tolerance mechanism. The *p*-value in panel C was obtained by a two-tailed, unpaired t-test of the fluorescence values. Blue lines and circles in panel A and bars in panels B and C show the geometric mean. Black dots indicate individual experiments. Errors are 95 % confidence intervals of the geometric mean obtained by bootstrapping. The grey area in panel A shows the detection limit. Experiments with zero-counts in panels A and B are indicated by triangles on the x-axis and were omitted.

We previously showed that a reduction in the net negative surface charge contributes to BAC tolerance in *E. coli* S4, presumably decreasing BAC adsorption to the cells ^12^. We corroborated these findings by utilizing the fluorescent properties of NBD-DDA to directly measure adsorption of NBD-DDA by the BAC susceptible wild-type and the BAC tolerant strain S4 (Figure 4 C). Due to its fluorescence, the accumulation of the antimicrobial NBD-DDA itself can be followed, preventing the need for an auxiliary fluorescent probe.

Taken together, our experiments indicate that the biological activity of NBD-DDA is very similar to that of BAC, not only in terms of its efficacy, but also in terms of mode-of-action. Furthermore, we showed that the tolerance against NBD-DDA, and its analogue BAC, is mediated through reduced adsorption by the tolerant *E. coli* strain S4.

### Fluorescence of NBD-DDA facilitates studies on the single-cell level

To demonstrate the potential of NBD-DDA for studying BAC and its mode-of-action on the molecular level and heterogeneity on the single-cell level, we stained the *E. coli* wild-type with NBD-DDA. To this end, the cells were incubated with a sub-inhibitory concentration of NBD-DDA (10 µM, 0.5 x MIC) and subsequently assessed by flow cytometry and confocal microscopy. *E. coli* cells stained with NBD-DDA were readily detected in the standard FITC/GFP channel (excitation: 488 nm, emission: 525 ± 20 nm) in the flow cytometer (Figure 5). The cells showed a fluorescence distribution, signifying the heterogeneity between individual cells (Figure 5 A). After 5 minutes of incubation, the population of fluorescently stained cells could be already clearly distinguished from the autofluorescence signal of unstained cells (Figure 5 A). A small fraction of around 9% of the cells incubated with NBD-DDA did not display a fluorescence signal different from that of the unlabeled cells (Figure 5 A). Increasing the staining duration increased the fluorescence signal derived from the bacteria cells (Figure 5 B), with half-maximal saturation of staining being reached after 5 minutes of incubation. The fraction of non-fluorescent cells was invariant to incubation times (data not shown).

**Figure 5:**
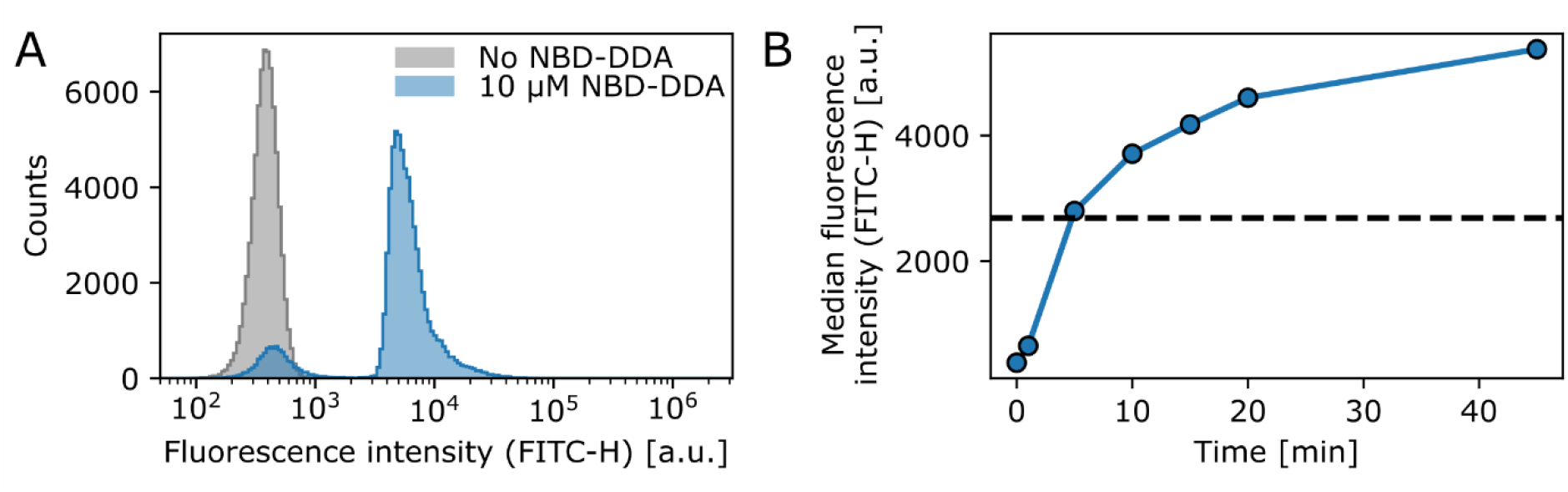
NBD-DDA effectively stains *E. coli* cells and can be detected with flow cytometry. A) Staining of *E. coli* wild-type with a sub inhibitory level of NBD-DDA. B) Staining with NBD-DDA increases with time, with the half maximal saturation of staining being reached after approx. 5 minutes (dashed line). The data points show the median fluorescence values of 50 000 – 70 000 cells. The fluorescence was recorded in the FITC channel (excitation at 488 nm; emission detection window of 525 ± 20 nm).). Given are the maximum intensity (height) values of the fluorescence signals.

*E. coli* cells stained with NBD-DDA were also examined using confocal microscopy (Figure 6 A). Fluorescence intensity was highest at the cell edges, suggesting that NBD-DDA is localized at the cell envelope. This observation was confirmed by co-staining of the cells with the membrane stain SynaptoRed C2. The signals from both fluorophores revealed dye colocalization (Figure 6 B and C). This is in accordance with the assumption that the cell membrane is the primary target of BAC ^2,3^. Thus, NBD-DDA could be leveraged in live cell imaging applications of mammalian and bacterial cells to label cell membranes ^21^.

**Figure 6:**
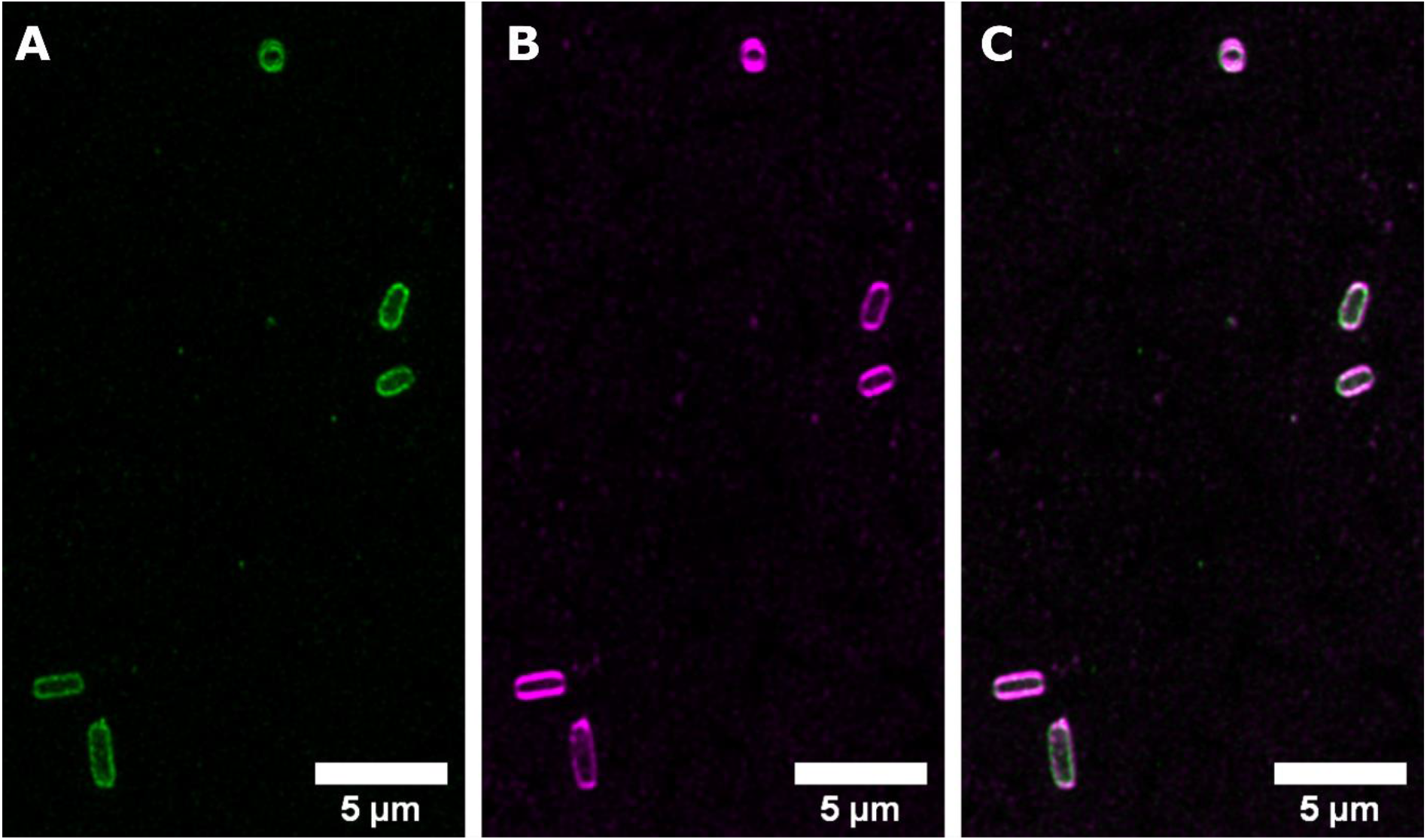
NBD-DDA is localized to the cell envelope of *E. coli*. *E. coli* cells stained with A) 10 µM NBD-DDA and B) with the membrane stain SynaptoRed C2 (10 µM). C) The overlay of both signals shows their colocalization. Shown are the maximum intensity projections of z-stacks with 10 slices. Image histograms were adjusted for visibility.

The potential of fluorescent probes derived from antimicrobial substances to study drug susceptibility in single bacterial cells has recently been demonstrated for a range of antibiotics ^22,23^. Here we demonstrate that this approach is also promising for BACs and related QACs, which are used as disinfectants, antiseptics, and preservatives ^1^. We used a sub-inhibitory concentration of NBD-DDA for staining purposes. In future studies, it will be interesting to observe the effects of higher concentrations on the localization within cells, as well as observing the dynamics of NBD-DDA adsorption and killing in individual cells, using time lapse microscopy in a microfluidic device.

## Conclusion and Outlook

We synthesized a fluorescent analogue of BAC, NBD-DDA that is readily detectable by fluorescence methods such as flow cytometry and fluorescence microscopy using standard FITC/GFP settings. Also, we showed that it can be utilized to study the mode-of-action and resistance mechanisms of bacteria against BAC and similar QACs, as well as a phenotypic heterogeneity on the single-cell level. NBD-DDA has three important properties:

- the antimicrobial activity of NBD-DDA is comparable to that of BAC,
- the mode of action of NBD-DDA is apparently similar to that of BAC,
- the fluorescent properties of NBD-DDA make it suitable to study phenotypic heterogeneity, efflux and cellular adsorption kinetics.

Leveraging these properties, we were able to gain three insights towards the mode of action of BAC:

- tolerance to NBD-DDA/BAC in *E. coli* is associated to reduced cellular adsorption
- adsorption of NBD-DDA/BAC to cells is heterogeneous within isogenic *E. coli* populations
- NBD-DDA/BAC adsorbs preferentially to the bacterial membrane

Our study highlights the potential and versatility of fluorescent QAC analogues, such as NBD-DDA. We anticipate that such compounds will facilitate the understanding of mechanisms related to disinfection, disinfection insusceptibility on the single-cell level, and cytotoxicity of disinfectants. For example, NBD-DDA could be used to:

- determine the mode of action of QACs against microorganisms.
- investigate the role of phenotypic heterogeneity on disinfection tolerance.
- identify efflux pumps which are specific to certain QACs. explore the mechanisms which underlie the geno- and cytotoxic effects of QACs in mammalian cells ^9,16–19^. Unraveling these mechanisms could guide appropriate counter measures to these effects in consumer products, such as ophthalmic solutions.
- investigate the distribution and fate of QACs when applied to surfaces and in environmental settings ^20^.

## Experimental Section

### Chemicals for synthesis of NBD-DDA

*N,N*-dimethylethylenediamine (Merck, 95%), 4-chloro-7-nitrobenzofurazan (NBD-Cl; Merck, 98%), 1-iodododecane (Merck, 98%), potassium acetate (Th. Geyer, 99%), ethanol (Th. Geyer, absolute, 99.9%), acetone (Th. Geyer, 99.5%), dichloromethane (DCM; Th. Geyer, HPLC-Grade, 99.9%), and methanol (Th. Geyer, 99.8%) were used as received without further purification. Spectroscopic measurements were carried out in spectroscopic grade dimethylsulfoxide (DMSO; Honeywell, absolute, for UV-spectroscopy, 99.8%) and phosphate buffered saline (PBS; 1x, pH = 7.4). PBS was prepared from a pre-weighed tablet (Th. Geyer) with Milli-Q water (> 18 MΩ) according to the instructions of the manufacturer.

### Spectrometry for product characterization

^1^H and ^13^C NMR spectra were recorded on an ECP500 (JEOL; NBD-DMA) or an ECP600 (JEOL; NBD-DDA) at a proton resonance frequency of 500 MHz and 600 MHz, respectively, and calibrated to the residual solvent signals at δ 7.26 (^1^H) and δ 77.16 (^13^C) for chloroform and δ 2.50 (^1^H) and δ 39.52 (^13^C) for DMSO ^24^, respectively. Electrospray ionization time-of-flight mass spectrometry ((ESI-TOF-MS) measurements were done with an Agilent 6210 ESI-TOF mass spectrometer (Agilent Technologies). The solvent flow rate was adjusted to 4 μl/min, the spray voltage set to 4 kV, and the drying gas flow rate set to 15 psi (1 bar). All other parameters were adjusted to yield a maximum abundance of the relative [M+H] peak. Fluorescence measurements were done at room temperature using dilute solutions of NBD-DDA in 10 mm x 10 mm quartz cuvettes (Hellma GmbH). Fluorescence emission measurements were performed with a calibrated spectrofluorometer FSP-920 (Edinburgh Photonics), equipped with a Xenon lamp and double monochromators. The fluorescence emission spectra were corrected for the wavelength dependent spectral responsivity of the emission channel (emission correction) and the fluorescence excitation spectra for the wavelength dependent spectral radiant power of the excitation spectra (excitation correction) ^25^.

### Synthesis of *N,N*-dimethyl-N’-(4-nitro-5-benzofurazanyl)-1,2-ethanediamine (NBD-DMA)

NBD-DMA was synthesized following a published procedure with slight modifications ^13^. In short, potassium acetate (1.083 g, 11 mmol) was added to a solution of *N,N*-dimethylethylenediamine (3 mL, 27 mmol) in ethanol (20 mL). NBD-Cl (1.1 g, 5.4 mmol) in ethanol (60 mL) was added slowly to this solution at 45°C and the reaction was stirred overnight. After removing volatiles *in vacuo*, the residue was taken up in water and extracted with ethyl acetate. Here, a much larger volume of solvents than described in the original publication had to be used during the extraction steps because of the strong coloration of both phases. The combined organic extracts were dried over magnesium sulfate and concentrated *in vacuo*. Purification by flash column chromatography on silica gel (70-230 mesh) with methanol:DCM = 6:94 (v/v) yielded the title compound as a brown solid (0.63 g, 2.5 mmol, 47%). ^1^H-NMR (CDCl_3_): δ 2.33 (s, 6H), δ 2.71 (t, 2H), δ 3.48 (m, 2H), δ 6.13 (d, 1H), δ 8.49 (d, 1H); ^13^C-NMR (CDCl_3_): δ 40.6, 45.1, 56.2, 98.9, 136.6, 144.1, 144.4; ESI-TOF-MS m/z: calculated for C_10_H_14_N_5_O_3_ ([M+H]^+^) 252.1091, observed 252.1153.

### Synthesis of *N*-dodecyl-*N,N*-dimethyl-[2-[(4-nitro-2,1,3-benzoxadiazol-7-yl)amino]ethyl]azanium-iodide (NBD-DDA)

0.3 g (1 mmol) of NBD-DMA was dissolved in 10 mL of acetone. 1-Iodododecane (0.36 ml, 5.08 mmol) was added, and the mixture was heated at 70°C until the reaction was complete. The amber-yellow ammonium salt was collected by vacuum filtration, washed 2x with acetone (2 × 10 ml), and dried at room temperature, yielding the title compound as an amber-yellow solid (0.246 g, 0.45 mmol, 45%). ^1^H-NMR (DMSO-d6): δ 0.84 (t, 3H), δ 1.24 (bm, 18H), δ 1.65 (m, 2H), δ 3.14 (s, 6H), δ 3.38 (m, 2H), δ 3.66 (t, 2H), δ 3.99 (m, 2H), δ 6.60 (d, 1H), δ 8.58 (d, 1H), δ 9.33 (s, 1H); ^13^C-NMR (DMSO-d6): δ 13.9, 21.8, 22.1, 25.7, 28.5, 28.7, 28.8, 28.9, 29.0, 31.3, 37.2, 50.6, 59.8, 62.1, 63.7, 100.1, 100.2, 122.1 137.6, 144.0, 144.5; ESI-TOF-MS m/z: calculated for C_22_H_38_N_5_O_3_ ([M-I]^+^) 420.2969, observed 420.3005.

### Bacterial strains and growth conditions

Bacteria were cultivated in 100 ml Erlenmeyer flasks filled with 10 ml medium at 37 °C with agitation at 220 rpm. *E. coli* K-12 MG1655, *S. aureus* SH1000 and *P. aeruginosa* MPAO1 ^26^ were used as reference strains to determine the antimicrobial activity of NBD-DDA. The BAC-tolerant strain *E. coli* S4 originates from an evolution experiment with BAC and *E. coli* K-12 MG1655 as ancestor ^12^. As previously described ^12^, *E. coli* were cultivated in M9 minimal medium containing: 42 mM Na_2_HPO_4_, 22 mM KH_2_PO_4_, 8.5 mM NaCl, 11.3 mM (NH_4_)_2_SO_4_, 1 mM MgSO_4_, 0.1 mM CaCl_2_, 0.2 mM Uracil, 1 µg/ml thiamine, trace elements (25 µM FeCl_3_, 4.95 µM ZnCl_2_, 2.1 µM CoCl_2_, 2 µM Na_2_MoO_4_, 1.7 µM CaCl_2_, 2.5 µM CuCl_2_, 2 µM H_3_BO_3_) and 20 mM glucose. Cultivation and susceptibility testing of *E. coli* was done in M9 to make the results obtained here comparable to the results we recently obtained for BAC ^12^. *S. aureus* and *P. aeruginosa* were cultivated in cation adjusted Mueller-Hinton broth 2 (MH2; Sigma-Aldrich, 90922). Plating for enumeration of colony forming units was done on LB agar medium (Lennox formulation; 10 g/l tryptone, 5 g/l yeast extract, 5 g/l NaCl, 15 g/l agar).

### Determination of MIC and MBC values

Stationary phase cultures were diluted to a final cell concentration of 10^7^ cfu/ml in a 96-well plate with wells holding a final reaction volume of 200 µl of fresh medium containing different concentrations of the investigated antimicrobial substance. The tested concentrations of BAC (Sigma-Aldrich, 234427), BAC_12_ (Sigma-Aldrich, 13380) and NBD-DDA were 10, 20, 30, 40, 60, 100, and 150 µM. The plates were incubated at 37 °C with agitation for 24 hours and growth was assessed visually. To determine the MBC, 10 µl from each well without visible growth were spotted onto LB agar medium and incubated at 37 °C for 24 hours. The first concentration that reduced the initial cell density by a factor of 10^3^ or more was designated the MBC. NBD-DDA was dissolved in DMSO. To assess the effect of DMSO on growth, a MIC assay with the solvent DMSO alone was conducted. Growth was not inhibited by DMSO in the concentration range tested here.

### Time-kill assays with *E. coli*

Time-kill assays were conducted as previously described ^12^. *E. coli* were inoculated into 10 ml M9 to a defined number of cells (10^4^ cfu/ml) and incubated at 37 °C with agitation at 220 rpm. After 24 hours, OD_600_ was determined and adjusted to an OD_600_ of 0.01 (∼10^7^ cfu/ml) in spent medium in 2 ml tubes (PP, LABSOLUTE) in a total volume of 900 µl. To maintain the cellular physiology in the stationary phase, time-kill assays were conducted in spent medium from the pre-culture, which was obtained by removing cells from the pre-culture medium by centrifugation for 2 minutes at 16000 g. After dilution of the cells, 10 µl were sampled to determine the initial cell concentration. Antimicrobials were added to the final concentrations indicated in Figure 4 and cultures were incubated at 37 °C with agitation (1200 rpm) in a tabletop Thermomixer (STARLAB). Samples of 10 µl were taken and immediately serially diluted in 96-well microplates with 90 µl phosphate buffered saline (PBS, pH 7; 10 mM Na_2_HPO_4_, 1.76 mM KH_2_PO_4_, 2.68 mM KCl, 137 mM NaCl) per well. To enumerate colony forming units (cfu), 10 µl from each well were spotted on square LB agar plates. Plates were left to dry and incubated at 30 °C for 16-24 hours. To determine the fraction of survivors for single time point incubations, the initial time point before addition of antimicrobial and the final time point after 20 minutes of exposure were sampled.

### Flow cytometry

A stationary phase culture of *E. coli* was diluted 1:200 in PBS filtered with a 0.2 µm syringe filter containing 10 µM of NBD-DDA. Cells were then incubated for different times and subjected to flow cytometry. Flow cytometric measurements were conducted using a Cytoflex S (Beckman-Coulter) with the following settings: flow rate 10 µl/min, primary threshold forward scatter-height (FSC) 1000, secondary threshold side scatter-height (SSC) 1000, and 30 s measurement time. These settings allowed the detection of unlabelled cells in the FSC and SSC channels. Between 50 000 – 70 000 events (i.e. cells) were recorded. NBD-DDA fluorescence was detected in the FITC channel using an excitation wavelength of 488 nm and an emission detection window of 525 ± 20 nm.

### Staining and confocal imaging

One ml of a stationary culture of *E. coli* were pelleted by centrifugation and resuspended in PBS. NBD-DDA and the membrane dye SynaptoRed C2 (Sigma-Aldrich, S6689) were added to a final concentration of 10 µM. The suspension was applied to an object slide and covered with a no. 1.5H cover slip (Marienfeld). Imaging was done on a confocal laser scanning microscope Leica SP8 equipped with a white light laser (Superk Extreme EXW-9 NIM, NKT Photonics, Denmark) using a 100x oil immersion objective with a numerical aperture of 1.4. The fluorescence of NBD-DDA was excited at 470 nm (laser power of 34%) and recorded in the spectral window of 500 – 600 nm. The membrane stain SynaptoRed C2 used for colocalization studies was excited at 570 nm (laser power 24.6%) and detected in the spectral window of 650 – 800 nm. Both channels were sequentially excited to avoid spectral bleed through. The pixel size of the recorded images was 41.4 nm.

A z-stack with 10 images and a step size of 0.26 µm was recorded. The images were deconvoluted with the software Huygens Essential (Version 17.04, Scientific Volume Imaging B.V., The Netherlands) with default settings and a maximum intensity projection of the deconvoluted z-stacks was created. The intensity histograms of the images were adjusted to improve the visibility of the fluorescence signal.

### NBD-DDA adsorption assay

Stationary phase cultures of *E. coli* were harvested by centrifugation for 4 minutes at 6000 g, the supernatant was removed and the OD_600_ was adjusted to 1 (approx. 10^9^ cfu/ml) in PBS and aliquoted into 200 µl samples. NBD-DDA was added to a final concentration of 10 µM and samples were incubated for 45 minutes. After another centrifugation step (4 min, 6000 g), the supernatant was removed, cells were resuspended in PBS and the fluorescence was determined with a Qubit 4 fluorometer (Thermo Fisher Scientific, Wilmington, DE, USA) with excitation at 470 nm and emission at 510 – 580 nm. Cells in PBS without NBD-DDA were used to determine the autofluorescence background, which was subtracted from the fluorescence signal obtained with NBD-DDA. Autofluorescence made up less than 1% of the fluorescence signal of the NBD-DDA treated samples.

## Acknowledgements

The authors acknowledge funding of the work by the Federal Institute for Materials Research and Testing (BAM). The work of KOH at BAM was performed as part of a traineeship funded by the European Union within the ERASMUS+ program.

## CRediT roles

**NN** Conceptualization, Data Curation, Formal Analysis, Investigation, Methodology, Software, Validation, Visualization, Writing – Original Draft Preparation, Writing – Review & Editing

**KOH** Formal Analysis, Funding Acquisition, Investigation, Writing – Review & Editing

**URG** Resources, Writing – Review & Editing

**MATB** Conceptualization, Writing – Review & Editing

**BR** Conceptualization, Data Curation, Formal Analysis, Investigation, Methodology, Supervision, Validation, Visualization, Writing – Original Draft Preparation, Writing – Review & Editing

**FS** Conceptualization, Formal Analysis, Funding Acquisition, Investigation, Methodology, Project Administration, Resources, Supervision, Visualization, Writing – Original Draft Preparation, Writing – Review & Editing

## References

(1) Pereira, B. M. P.; Tagkopoulos, I. Benzalkonium Chlorides: Uses, Regulatory Status, and Microbial Resistance. Appl. Environ. Microbiol. 2019, 85 (13). https://doi.org/10.1128/AEM.00377-19.

(2) Gilbert, P.; Moore, L. E. Cationic Antiseptics: Diversity of Action under a Common Epithet. Journal of Applied Microbiology 2005, 99 (4), 703–715. https://doi.org/10.1111/j.1365-2672.2005.02664.x.

(3) Kwaśniewska, D.; Chen, Y.-L.; Wieczorek, D. Biological Activity of Quaternary Ammonium Salts and Their Derivatives. Pathogens 2020, 9 (6), 459. https://doi.org/10.3390/pathogens9060459.

(4) Jono, K.; Takayama, T.; Kuno, M.; Higashide, E. Effect of Alkyl Chain Length of Benzalkonium Chloride on the Bactericidal Activity and Binding to Organic Materials. Chemical & Pharmaceutical Bulletin 1986, 34 (10), 4215–4224. https://doi.org/10.1248/cpb.34.4215.

(5) Daoud, N. N.; Dickinson, N. A.; Gilbert, P. Antimicrobial Activity and Physico-Chemical Properties of Some Alkyldimethylbenzylammonium Chlorides. Microbios 1983, 37 (148), 73–85.

(6) DeLeo, P. C.; Huynh, C.; Pattanayek, M.; Schmid, K. C.; Pechacek, N. Assessment of Ecological Hazards and Environmental Fate of Disinfectant Quaternary Ammonium Compounds. Ecotoxicol Environ Saf 2020, 206, 111116. https://doi.org/10.1016/j.ecoenv.2020.111116.

(7) Barros, A. C.; Melo, L. F.; Pereira, A. A Multi-Purpose Approach to the Mechanisms of Action of Two Biocides (Benzalkonium Chloride and Dibromonitrilopropionamide): Discussion of Pseudomonas Fluorescens’ Viability and Death. Frontiers in Microbiology 2022, 13.

(8) Tischer, M.; Pradel, G.; Ohlsen, K.; Holzgrabe, U. Quaternary Ammonium Salts and Their Antimicrobial Potential: Targets or Nonspecific Interactions? ChemMedChem 2012, 7 (1), 22–31. https://doi.org/10.1002/cmdc.201100404.

(9) Zhang, S.; Ding, S.; Yu, J.; Chen, X.; Lei, Q.; Fang, W. Antibacterial Activity, in Vitro Cytotoxicity, and Cell Cycle Arrest of Gemini Quaternary Ammonium Surfactants. Langmuir 2015, 31 (44), 12161–12169. https://doi.org/10.1021/acs.langmuir.5b01430.

(10) Inácio, Â. S.; Domingues, N. S.; Nunes, A.; Martins, P. T.; Moreno, M. J.; Estronca, L. M.; Fernandes, R.; Moreno, A. J. M.; Borrego, M. J.; Gomes, J. P.; Vaz, W. L. C.; Vieira, O. V. Quaternary Ammonium Surfactant Structure Determines Selective Toxicity towards Bacteria: Mechanisms of Action and Clinical Implications in Antibacterial Prophylaxis. Journal of Antimicrobial Chemotherapy 2016, 71 (3), 641–654. https://doi.org/10.1093/jac/dkv405.

(11) Kampf, G. Antiseptic Stewardship: Biocide Resistance and Clinical Implications; Springer International Publishing, 2018. https://doi.org/10.1007/978-3-319-98785-9.

(12) Nordholt, N.; Kanaris, O.; Schmidt, S. B. I.; Schreiber, F. Persistence against Benzalkonium Chloride Promotes Rapid Evolution of Tolerance during Periodic Disinfection. Nat Commun 2021, 12 (1), 6792. https://doi.org/10.1038/s41467-021-27019-8.

(13) Bednarczyk, D.; Mash, E. A.; Aavula, B. R.; Wright, S. H. NBD-TMA: A Novel Fluorescent Substrate of the Peritubular Organic Cation Transporter of Renal Proximal Tubules. Eur J Physiol 2000, 440 (1), 184–192. https://doi.org/10.1007/s004240000283.

(14) Amaro, M.; Filipe, H. A. L.; Ramalho, J. P. P.; Hof, M.; Loura, L. M. S. Fluorescence of Nitrobenzoxadiazole (NBD)-Labeled Lipids in Model Membranes Is Connected Not to Lipid Mobility but to Probe Location. Phys. Chem. Chem. Phys. 2016, 18 (10), 7042–7054. https://doi.org/10.1039/C5CP05238F.

(15) Chattopadhyay, A. Chemistry and Biology of N-(7-Nitrobenz-2-Oxa-1,3-Diazol-4-Yl)-Labeled Lipids: Fluorescent Probes of Biological and Model Membranes. Chemistry and Physics of Lipids 1990, 53 (1), 1–15. https://doi.org/10.1016/0009-3084(90)90128-E.

(16) De Saint Jean, M.; Brignole, F.; Bringuier, A. F.; Bauchet, A.; Feldmann, G.; Baudouin, C. Effects of Benzalkonium Chloride on Growth and Survival of Chang Conjunctival Cells. Invest Ophthalmol Vis Sci 1999, 40 (3), 619–630.

(17) Ferk, F.; Misík, M.; Hoelzl, C.; Uhl, M.; Fuerhacker, M.; Grillitsch, B.; Parzefall, W.; Nersesyan, A.; Micieta, K.; Grummt, T.; Ehrlich, V.; Knasmüller, S. Benzalkonium Chloride (BAC) and Dimethyldioctadecyl-Ammonium Bromide (DDAB), Two Common Quaternary Ammonium Compounds, Cause Genotoxic Effects in Mammalian and Plant Cells at Environmentally Relevant Concentrations. Mutagenesis 2007, 22 (6), 363–370. https://doi.org/10.1093/mutage/gem027.

(18) Datta, S.; Baudouin, C.; Brignole-Baudouin, F.; Denoyer, A.; Cortopassi, G. A. The Eye Drop Preservative Benzalkonium Chloride Potently Induces Mitochondrial Dysfunction and Preferentially Affects LHON Mutant Cells. Invest Ophthalmol Vis Sci 2017, 58 (4), 2406–2412. https://doi.org/10.1167/iovs.16-20903.

(19) Goldstein, M. H.; Silva, F. Q.; Blender, N.; Tran, T.; Vantipalli, S. Ocular Benzalkonium Chloride Exposure: Problems and Solutions. Eye 2022, 36 (2), 361–368. https://doi.org/10.1038/s41433-021-01668-x.

(20) Wood, J. P.; Magnuson, M.; Touati, A.; Gilberry, J.; Sawyer, J.; Chamberlain, T.; McDonald, S.; Hook, D. Evaluation of Electrostatic Sprayers and Foggers for the Application of Disinfectants in the Era of SARS-CoV-2. PLoS One 2021, 16 (9), e0257434. https://doi.org/10.1371/journal.pone.0257434.

(21) Deng, P.; Xiao, F.; Wang, Z.; Jin, G. A Novel BODIPY Quaternary Ammonium Salt-Based Fluorescent Probe: Synthesis, Physical Properties, and Live-Cell Imaging. Frontiers in Chemistry 2021, 9.

(22) Łapińska, U.; Voliotis, M.; Lee, K. K.; Campey, A.; Stone, M. R. L.; Tuck, B.; Phetsang, W.; Zhang, B.; Tsaneva-Atanasova, K.; Blaskovich, M. A.; Pagliara, S. Fast Bacterial Growth Reduces Antibiotic Accumulation and Efficacy. eLife 2022, 11, e74062. https://doi.org/10.7554/eLife.74062.

(23) Stone, M. R. L.; Masi, M.; Phetsang, W.; Pagès, J.-M.; Cooper, M. A.; Blaskovich, M. A. T. Fluoroquinolone-Derived Fluorescent Probes for Studies of Bacterial Penetration and Efflux. Med. Chem. Commun. 2019, 10 (6), 901–906. https://doi.org/10.1039/C9MD00124G.

(24) Fulmer, G. R.; Miller, A. J. M.; Sherden, N. H.; Gottlieb, H. E.; Nudelman, A.; Stoltz, B. M.; Bercaw, J. E.; Goldberg, K. I. NMR Chemical Shifts of Trace Impurities: Common Laboratory Solvents, Organics, and Gases in Deuterated Solvents Relevant to the Organometallic Chemist. Organometallics 2010, 29 (9), 2176–2179. https://doi.org/10.1021/om100106e.

(25) Resch-Genger, U.; Pfeifer, D.; Monte, C.; Pilz, W.; Hoffmann, A.; Spieles, M.; Rurack, K.; Hollandt, J.; Taubert, D.; Schönenberger, B.; Nording, P. Traceability in Fluorometry: Part II. Spectral Fluorescence Standards. J Fluoresc 2005, 15 (3), 315–336. https://doi.org/10.1007/s10895-005-2629-9.

(26) Varadarajan, A. R.; Allan, R. N.; Valentin, J. D. P.; Castañeda Ocampo, O. E.; Somerville, V.; Pietsch, F.; Buhmann, M. T.; West, J.; Skipp, P. J.; van der Mei, H. C.; Ren, Q.; Schreiber, F.; Webb, J. S.; Ahrens, C. H. An Integrated Model System to Gain Mechanistic Insights into Biofilm-Associated Antimicrobial Resistance in Pseudomonas Aeruginosa MPAO1. npj Biofilms and Microbiomes 2020, 6 (1), 1–17. https://doi.org/10.1038/s41522-020-00154-8.

